# A novel PLS1 c.981+1G>A variant causes autosomal-dominant hereditary hearing loss in a family via up-regulation of the PI3K-Akt signaling pathway

**DOI:** 10.1101/2022.03.17.484618

**Authors:** Liangpu Xu, Xinrui Wang, Jia Li, Lingji Chen, Haiwei Wang, Shiyi Xu, Yanhong Zhang, Wei Li, Pengcheng Yao, Meihua Tan, Si Zhou, Meihuan Chen, Yali Pan, Xuemei Chen, Xiaolan Chen, Yunliang Liu, Na Lin, Hailong Huang, Hua Cao

**Author notes:** Co-first authors:Liangpu Xu,Xinrui Wang,Jia Li,Lingji Chen. Correspondence author: Liangpu Xu,; Na Lin,; Hailong Huang,;Hua Cao.

## Abstract

**Background:** The fimbrin protein family contains a variety of proteins, of which Plastin1 (PLS1) is an important member. The latest researches show that variations in the coding region of PLS1 gene is related to the development of deafness.

However, it remains unknown about the molecular mechanism of deafness caused by *PLS1* gene variants.

**Methods:** whole exome sequencing was performed on the affected family members and healthy ones to identify the pathogenic variants, followed by sanger sequencing. Minigene assay was conducted to investigate the impact of the variant on *PLS1* mRNA splicing. The pathogenicity of the variant was further investigated in zebrafish. RNA-sequencing (RNA-seq) was done for analyzing the dysregulated downstream signaling pathways caused by knockdown of *PLS1* expression.

**Results:** We identified a novel variant *PLS1* c.981+1G>A in a large Chinese family with hearing loss and demonstrated that the variant is responsible for the occurrence of hearing loss by inducing Exon8 skipping or deletion of Exon8 (47bp). The variant caused abnormal inner ear phenotypes, characterized by decreased mean otolith distance, anterior otolith diameter, posterior otolith diameter, and cochlear diameter, swimming speed and distance in zebrafish. Furthermore, silencing *PLS1* expression significantly upregulated genes in the PI3K-Akt signaling pathway, including Col6a3, Spp1, Itgb3 and Hgf.

**Conclusions:** *PLS1* c.981+1G>A is a novel pathogenic variant for hearing loss by inducing Exon8 skipping or deletion of Exon8 (47bp). Up-regulation of the PI3K-Akt signaling pathway plays an important role in the pathogenesis of PLS-1 gene.

## Introduction

Hearing loss is one of the most common sensorial diseases, affecting approximately 1 in 1,000 children Genetic factors account for at least half of the cases^1^. Among them, 30% is syndromic hearing loss (SHL) and the rest is non-syndromic hearing loss (NSHL) which is absent of additional symptoms. The majority of NSHL are Mendelian monogenetic disorders. Till now, more than one hundred NSHL genes have been defined. However, because the high heterogeneity of hearing loss, there are many patients and families with hereditary hearing loss whose causative genes and variants have not been identified^2^. Through Whole-exome sequencing (WES), a large number of mutations causing hearing loss has been successfully identified.

By using NGS, a large number of new hearing loss causing genes has been defined, such as LALYlA^3^, COLllAl^4^, PDElC^5^ and IFNLRI ^6^.

Plastin1 (PLS1) is an important member of the fimbrin protein family. It was first found in small intestine, colon and kidney^7^. Silencing PLS1 gene expression in mice has abnormal ultrastructural changes of small intestine, including shortening of small intestinal microvilli and contraction of base. Its core actin bundle obviously lacks real foundation, resulting in increased vulnerability of intestinal epithelium and reduced cross epithelial resistance, indicating that PLS1 is an important regulator of the morphology and stability of intestinal villous brush border^8^. PLS1 drives colorectal cancer metastasis by regulating IQGAP1/Rac1/ERK signaling pathway^9^. PLS1 promotes osteoblast differentiation by regulating intracellular calcium concentration^10^. In rat cochlear hair cells, PLS1 protein is continuously expressed from early stage to adult stage, indicating that PLS1 protein is the main component of adult cochlear hair cells. The structure of PLS1 protein cross-linked actin filament may be the key for cochlear hair cells to transmit auditory signals^11^. Further studies found that silencing PLS1 gene expression in mice could lead to abnormal maintenance of stereocilia of mouse hair cells and moderate hearing loss^12,13^. In recent years, some researchers have reported that the variation in the coding region of *PLS1* gene is related to the occurrence of deafness^14-16^. However, it remains unknown about the molecular mechanism of deafness caused by *PLS1* gene mutations.

In the present study, using WES we identified a novel variant *PLS1* c.981+1G>A in a large Chinese family with hearing loss. The pathogenesis of abnormal mRNA splicing induced by a novel variant of *PLS1* c.981+1G>A has been analyzed. We demonstrate that the variant is responsible for the occurrence of hearing loss by inducing aberrant splicing pattern of *PLS1* mRNA in the family. The variant caused abnormal inner ear phenotypes, characterized by decreased mean otolith distance, anterior otolith diameter, posterior otolith diameter, cochlear diameter in zebrafish, reduced swimming speed and distance. Further analysis revealed *PLS-1* is implicated in hearing loss probably via activating PI3K-Akt signaling pathway.

## Methods and materials

### Patients and clinical evaluations

The hearing loss family has 12 affected members and 29 unaffected members. Venous blood samples (5 ml) were obtained from 12 affected members and 5 healthy members. Pure-tone hearing testing was done for 17 family members, and we measured air conduction thresholds at 250 Hz, 500 Hz, 1 kHz, 2 kHz, 4 kHz, 6 kHz, and 8 kHz. We also determined bone conduction thresholds to exclude conductive components in patients with hearing loss. Written informed consent was obtained from all participants. Our research followed the Declaration of Helsinki. It has been approved by the Institutional Review Board of Fujian Maternal and Child Health Hospital (No. 2020YJ223-02) and Beijing Genome Research Institute (BGI-IRB 21129)

### Library preparation and next-generation sequencing

Genomic DNA was extracted from 5ml venous blood of the 17 family members following the manufacturer’s protocol of the MagPure Buffy Coat DNA Midi KF Kit (Magen, Guangzhou, China) and then fragmented by Segmentase (BGI, China). The fragments with size ranging from 280bp to 320bp were further enriched by magnetic bead. After end-filling and addition of base “A”, the ligation-mediated polymerase chain reaction was used to amplify DNA fragments, and finally purify to form a library. Following the manufacturer’s protocol (Roche nimblegen, Madison, USA), the library was enriched by array hybridization, and then eluted and post-capture amplified. Then, the enrichment degree of the library was measured on Agilent 2100 Bioanalyzer. MGIseq-2000 platform was used to sequence all qualified libraries with a sequencing strategy of Paired-end 100

### Bioinformatics analysis and validation of pathogenic variant

We used previously published filtering criteria to generate “clean reads” for downstream analysis ^17^. The “clean reads” were then aligned against the human genome reference 19 (hg19) using the Burrows Wheeler Aligner (BWA) software^18^. After alignment and removal of duplicated reads, GATK V3.6 was used to detect single-nucleotide variants (SNVs) and insertions and deletions (indels). The minor allele frequencies (MAF) of all SNVs and indels were annotated using multiple databases, including dbSNP^19^, HapMap^20^, The Exome Aggregation Consortium^21^(ExAC), The Genome Aggregation Database^22^ (gnomAD), 1000 human genome dataset^23^ and database of 100 Chinese healthy adults. Variants with MAF < 0.01 were eliminated from further analysis. The functional effects of variants were evaluated by The Sorting Intolerant from Tolerant (SIFT) and Polyphen2. The evaluation of pathogenic variants was carried out in accordance with the guidelines issued by American College of Medical Genetics (ACMG)^24^. The Human Gene Mutation Database^25^ (HGMD) was used to identify known pathogenic variants. *PLS1* c.981+1G>A (NM_001172312) was validated for the 12 affected members and 5 healthy family members using the conventional Sanger sequencing.

### Prediction of splicing effect, model building and structural-based analysis

We obtained wildtype and mutant sequences in FASTA format from NCBI GenBank (NM_001172312) and utilized augutus to predict the effect of *PLS1* c.981+1G>A on alternative splicing (https://bioinf.uni-greifswald.de/augustus/submission.php). Then, the wildtype protein sequence of human PLS1, including its 629 amino acids and its mutation were accessed from the augutus website and subjected to SWISS-MODEL^26^ (http://swissmodel.expasy.org/workspace/) for three dimensional modeling. Last, the software PyMOL was used to visualize and compare the structures of wildtype and mutant PLS1 proteins.

### Cell culture conditions

The HEI-OC1 cell line was grown in DMEM (GibcoBRL, Gaithersburg, MD, USA), supplemented with 10% FBS (GibcoBRL, Gaithersburg, MD, USA) at 33°C in a humidified atmosphere containing 10% CO_2_. The Human MCF-7 and 293T cells were maintained in DMEM supplemented with 10% fetal bovine serum at 37°C and 5% CO_2_.

### Minigene splicing assays

We performed minigene splicing assays based on the comparative analysis of the splicing pattern of wild-type (WT) and mutant reporter minigenes to evaluate the impact of *PLS1* c.981+1G>A on mRNA splicing. We prepared minigenes using the pcMINI and pcMINI-N vector backbone and cloned DNA fragments containing surrounding exons and introns of the c.981+1G>A variant in the PLS1 gene using classical restriction and ligation methods. After incubation for 36 h, we used RNeasy Mini Kit (Qiagen, Germany) to extract total RNA, and then reverse transcribed RNA to get cDNA using PrimeScript RT reagent kit (Takara, Osaka, Japan). Supplementary Table 1 lists the primers used to construct minigene vectors and to detect alternative splice sites.

### shRNA□induced gene silencing by lentiviral infection

Lentivirus packaging was made by GenePharma Inc (Shanghai, China). Lentiviral shRNAs targeting Pls1 were obtained from Genepharma, containing shRNAs scrambled (against the sequence 5[-TTCTCCGAACGTGTCACGT-3[), and Pls1 (against the sequence 5[-GCTCCAGGAAGCGGATAAACT-3[and 5[-GCCATTGATCTTTCAGGATTT-3[). HEI-OC1 cells were cultured with virus containing medium and Polybrene at a final concentration of 10 μg/mL. 0.5 μg/mL puromycin was used to screen infected cells.RNA-sequencing (RNA-seq) was performed to investigate the downstream dysregulated signaling pathway after knockdown of Pls-1 in HEI-OC1 cells. Detailed information regarding the library preparation and bioinformatic analysis of the RNA-seq analysis was presented in Supplementary Method 1.

### Western blotting

We washed cells twice with pre-chilled PBS and then lysed cells with radioimmunoprecipitation buffer (RIPA). The lysed cells were transferred to a centrifuge tube, centrifuged at 14,000 rpm for 10 min at 4°C (pre-cooling the centrifuge in advance). We recovered supernatants and quantified the protein by Bradford assay.All protein samples (10 μg) were electrophoresed on 10% SDS-PAGE gels, transferred to polyvinylidene difluoride membranes, and blocked in 5% skim milk in Tris Buffered Saline with Tween 20 for 2 h. Membranes were incubated with primary antibody at 4°C overnight, followed by incubation with horseradish peroxidase conjugated-secondary antibody for 2 h. The following primary antibodies were used in this study: anti-PLS1 (A15303, ABclonal, Wuhan, China), anti β-actin (Cat#5125, Cell Signaling Technology, Danvers, MA, USA).

### Preparation and microinjection of mRNA samples

The zebrafish experiments were carried out under the comprehensive animal care and use guidelines. The wild-type line AB was used in this study. Human mutant and wildtype pls1 cDNAs were synthesized (ShangYa Company, China) and subcloned into the Pmax vector. Mutant and wildtype pls1 open reading frames (ORF) were amplified from Pmax vector with SP6 promoter (Forward Primer:5’-CATACGATTTAGGTGACACTATAGCCGCCACCATGGA AAACAGTACTACTACCATTTCT-3’; Reverse Primer: 5’-CCAGATCTCGAGCT CGATGA-3’). We utilized mMESSAGE mMACHINE SP6 Transcription Kit (Ambion, USA, AM1340) to perform transcriptions and added Poly(A) signal to the 3’ end of capped mRNAs using Poly(A) Tailing Kit (Ambion, USA, AM1350).1-2 nL (50 ng/μL) solution containing capped mRNAs (green fluorescence protein [GFP]mRNA or mutant pls1 mRNA or wildtype pls1 mRNA) was injected into the zebrafish embryos (one-cell stage) using FemtoJet 4i (Eppendorf, Germany). Embryos at different developmental stages were preserved and collected for subsequent experiments

### Measurement of inner ears of Zebrafish

72 hours post-fertilization (hpf) zebrafish microinjected with different kinds of mRNAs were collected and photographed in a standardized lateral positioning. These images were taken under a stereo fluorescence microscope with a 30x magnification. Otolith distance, anterior otolith diameter, posterior otolith diameter and cochlear diameter were measured by stereo fluorescence microscope (Nikon SM2745T, Japan).

### Zebrafish behavioral assay

In order to observe the behavior trajectory of mutant pls1 overexpression and wildtype pls1 overexpression zebrafish, 6 days post-fertilization (dpf) larvae were placed into 96-well cell culture plates individually with 0.2 mL ddH_2_0. After standing for 10 min, the larvae were carefully placed into the instrument. Videos were obtained by BASLER aca-1300-60GM video tracking software (BASLER, Germany). 48 zebrafish of each group were tracked and recorded in a closed environment with 85 dB noise. Total swimming distance and speed were measured using the software zebrazoom^27^ (https://zebrazoom.org/).

### Statistical analysis

All data were expressed as mean + standard deviation and analyzed with Prism 5(GraphPad Software Company, America). Quantitative variables between groups were compared using Student t-test. P < 0.05 was considered statistically significant.

## Results

### Identification of a hearing loss family

We identified a hearing-loss family in Fujian province, the proband (III-8) presented congenital hearing loss and needed to wear hearing aids. The hearing loss family has 12 affected members and 29 unaffected members (Figure1A). Among the hearing loss patients in the family, the majority of patients presented bilateral severe-extremely severe sensorineural hearing loss except for II-8, III-12, III-15 and IV-8, which were bilateral moderate-severe sensorineural hearing loss. Pure-tone hearing testing showed the patient’s sensorineural hearing loss, and the audiogram shows a flat or declining curve (Figure1B-E).

**Figure 1.**
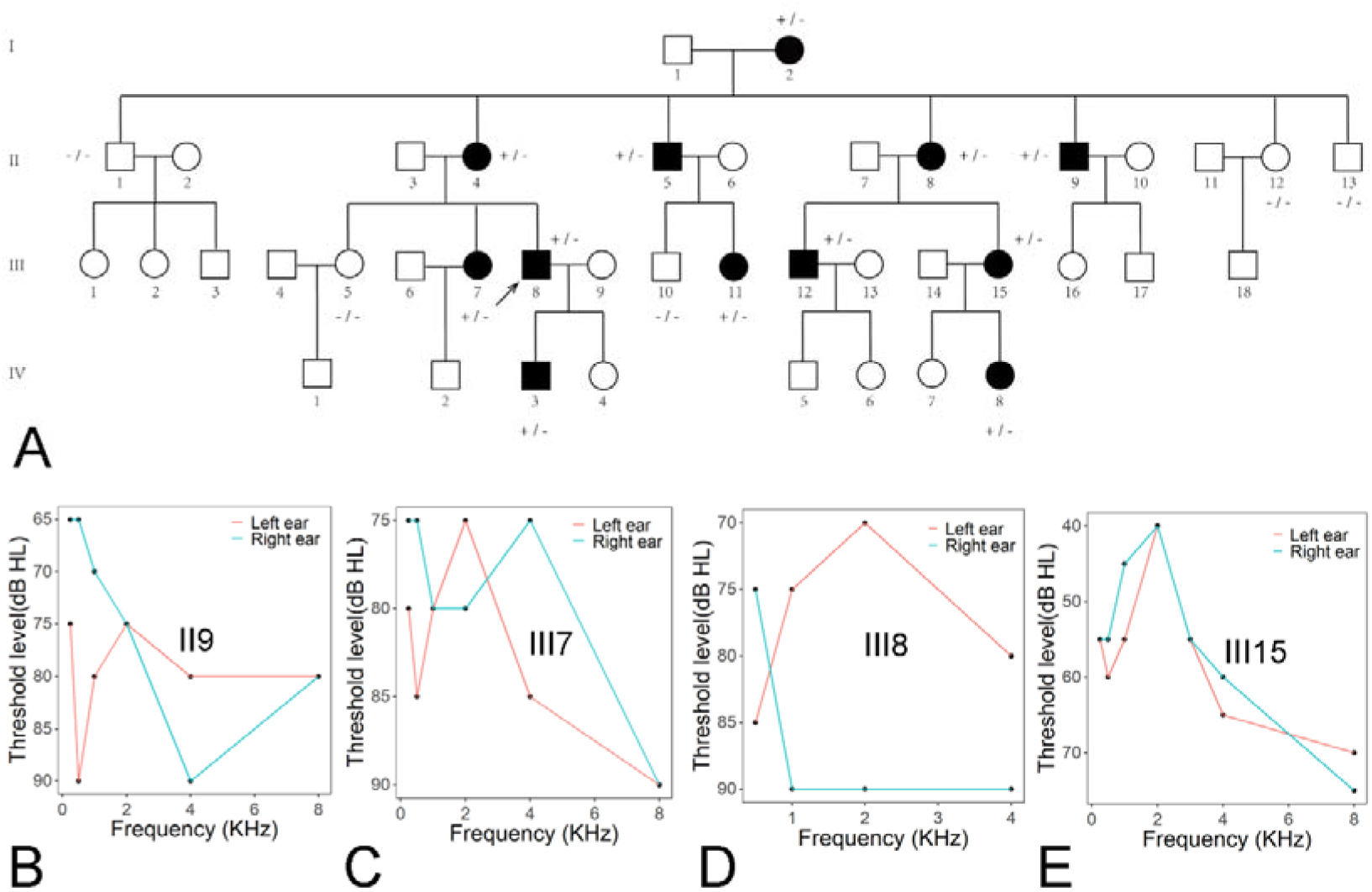
Identification of a large family with hearing loss. A. Pedigree of the hearing loss family, the arrow indicates the proband. Affected and unaffected family members are shown in black and white respectively. +/- and -/- indicate family members who are heterogeneous or wildtype for the *PLS1* c.981+1G>A variant. Pure-tone hearing testing results show flat or declining curves in the family members II9 (B), III7(C), III8 (D) and III15 (E).

### Exome sequencing identifies *PLS1* c.981+1G>A as likely pathogenic variant

To characterize the genetic cause of the hearing loss family, we sequenced the exome of the 12 affected members and 5 healthy family members in the hearing loss family. 20.67 billion bases of sequence were generated and aligned to the human reference 19. The total reads were mapped to the targeted bases with a coverage depth of 235.21-fold and coverage rate of 99.79%. We detected a total of 623,139 SNVs and indels in 17 family members.. SNVs and indels were filtered by dbSNP129, HapMap8, EXAC, GnomAD and the 1000 genome project and the shared SNPs were removed.

We used SIFT and PolyPhen2 to assess the non-synonymous variants for a likely functional impact. As compared to the Online Mendelian Inheritance in Man (OMIM) database^28^, we found a suspiciously pathogenic heterogeneous variant, c.981+1G>A (NM_001172312) in the splicing sites of *PLS1*. The variant was co-segregated with the hearing loss phenotype in the family. Sanger sequencing confirmed the heterogeneous *PLS1* c.981+1G>A variant was present in all hearing loss patients but absent in 5 healthy individuals (Supplementary Figure1).

### *PLS1* c.981+1G>A variant leads to Exon8 skipping or deletion of Exon8 (47bp)

As the *PLS1* c.981+1G>A variant is a splicing variant, we first utilized the online software augutus to predict the impact of the variant on the splicing pattern of *PLS1*. The results showed the variant led to Exon8 skipping (Supplementary Figure2). With the SWISS-MODEL method, we show the *PLS1* c.981+1G>A variant disrupted mRNA splicing and skipping of Exon 8 (amino acid: 297-327) and produced a shorted abnormal protein (Supplementary Figure3). Thus, we speculated that the *PLS1* c.981+1G>A variant was responsible for the hearing loss phenotype by inducing aberrant deletion of exon 8 and aberrant translation of PLS1 protein in the family.

To further validate our hypothesis, we performed minigene assay to investigate the splicing effects of *PLS1* c.981+1G>A variant. The entire Exon8(93bp) and surrounding intron 7(344bp) and intron 8(384bp) of wild-type and mutant PLS1 were cloned into a report vector pcMINI and transfected into HEK293T and MCF7 human cell lines (Figure2A). Urea-PAGE of reverse transcription polymerase chain reaction (RT-PCR) fragments generated from minigene spliced RNA of wild type and mutant constructions showed that the wildtype construction (pcMINI) presented an expected size of band, namely band a (482bp). The mutant construction presented a smaller DNA band, namely band b in HEK293T and MCF7 cell lines (Figure 2B). Sanger sequencing showed the wild type band had a normal splicing pattern [ExonA-Exon8-ExonB], while, the band b contained partial mRNA (ExonA-ExonB) and exon 8 skipping (Figure 2C). The entire Exon8(93bp) and intron 8(798bp) of wild-type and mutant PLS1 were cloned into the pcMINI vector. Minigene assay confirmed *PLS1* c.981+1G>A variant led to partial mRNA expression and there was an 47bp deletion on the right side of Exon8 (Supplementary Figure4). Therefore, *PLS1* c.981+1G>A variant leads to the abnormal splicing pattern of mRNA, resulting in complete skipping of Exon8 or deletion of partial Exon8 (47bp).

**Figure 2.**
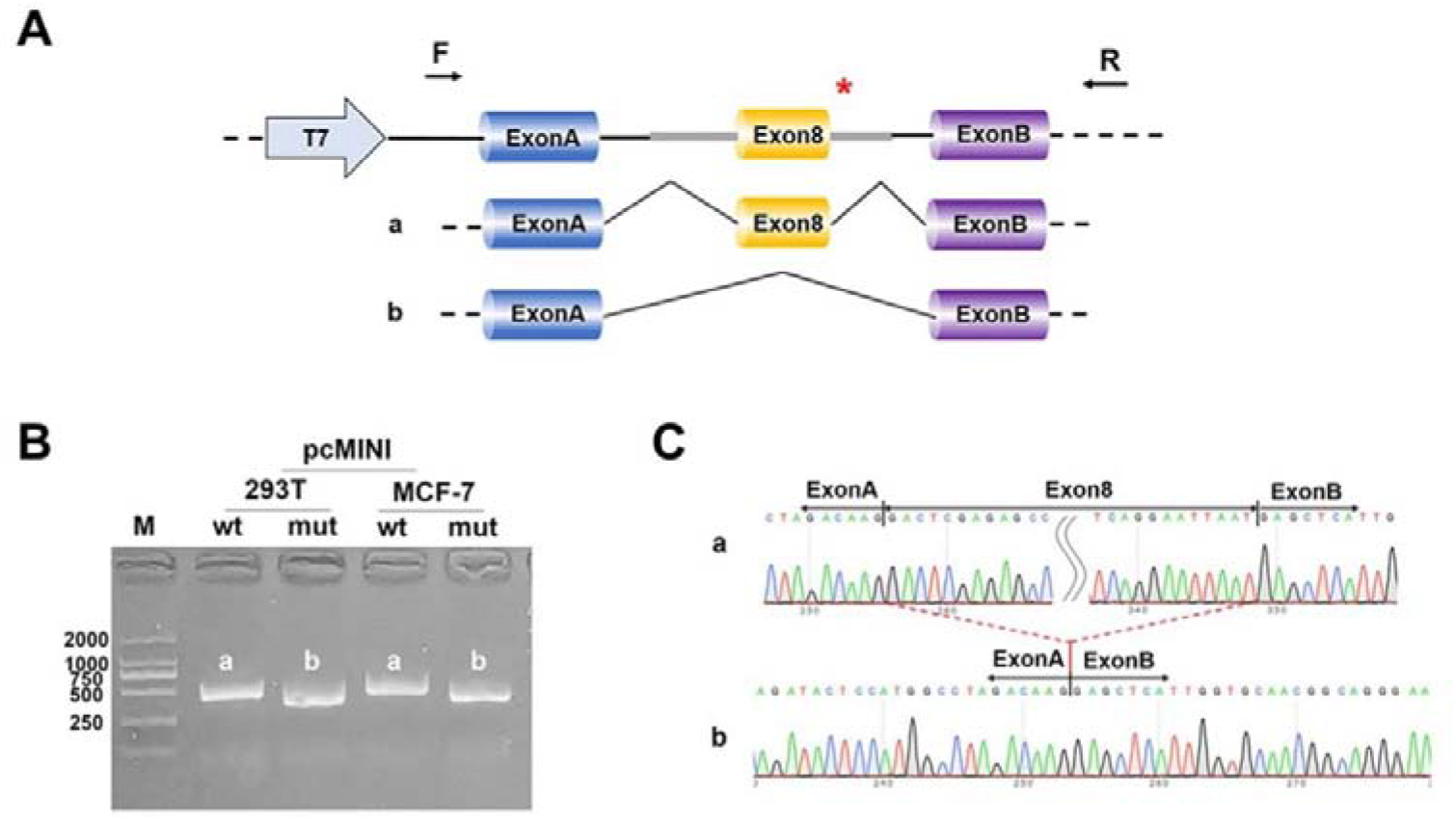
Results of minigene assay for *PLS1* c.981+1G>A in the HEK293T and MCF7 cell lines. A: The entire Exon8 (93bp) and surrounding intron 7(344bp) and intron 8(384bp) of wild-type and mutant *PLS1* were cloned into a report vector, pcMINI and transfected into HEK293T and MCF7 human cell lines. B. RT-PCR results of pcMINI-PLS1-mut and pcMINI-PLS1-wt constructs showed different sizes of bands in HEK293T and MCF-7 cells; C. Sanger sequencing shows that band a presented a normal splicing pattern, while band b had skipping of exon8.

### *PLS1* c.981+1G>A variant caused abnormal inner ear phenotypes in Zebrafish

2 nL solution containing capped mRNA sample (GFP mRNA as negative control or mutant pls1 mRNA or wildtype pls1 mRNA) was injected into the zebrafish embryos. After 24 hours, GFP was expressed in the zebrafish embryos. By observing the fluorescence expression, the embryos with GFP mRNA, mutant pls1 mRNA and wildtype pls1 mRNA expression were selected for follow-up experiments (Supplementary Figure5). In this study, 72 hpf zebrafish microinjected with different kinds of mRNAs were collected and photographed in a standardized lateral positioning. Otolith distance, anterior otolith diameter, posterior otolith diameter, and cochlear diameter were measured and calculated. The otolith distances were 0.02 ± 0.01 mm, 0.07 ± 0.01 mm, and 0.01 ± 0.01 mm in GFP, wildtype pls1, mutant pls1 groups, respectively. The mean otolith distance was shorter in mutant pls1 than that of the wildtype pls1 group (P< 0.05, Unpaired t test with Welch’s correction, Figure3). Moreover, anterior otolith diameter, posterior otolith diameter, and cochlear diameter were significantly decreased in mutant pls1 group than those in wildtype pls1 group (P< 0.05 for all cases, Unpaired t test with Welch’s correction, Figure3). Furthermore, 96 wildtype pls1 and mutant pls1 larvae were tracked with BASLER aca-1300-60GM video tracking software. Total swimming distance and speed were measured using the software zebrazoom. The average swimming distance and speed were calculated. The wildtype pls1 zebrafish exhibited significantly longer mean swimming distance and faster swimming speed than the mutant pls1 zebrafish (P< 0.05 for all cases, Unpaired t test with Welch’s correction, Figure4).

**Figure 3.**
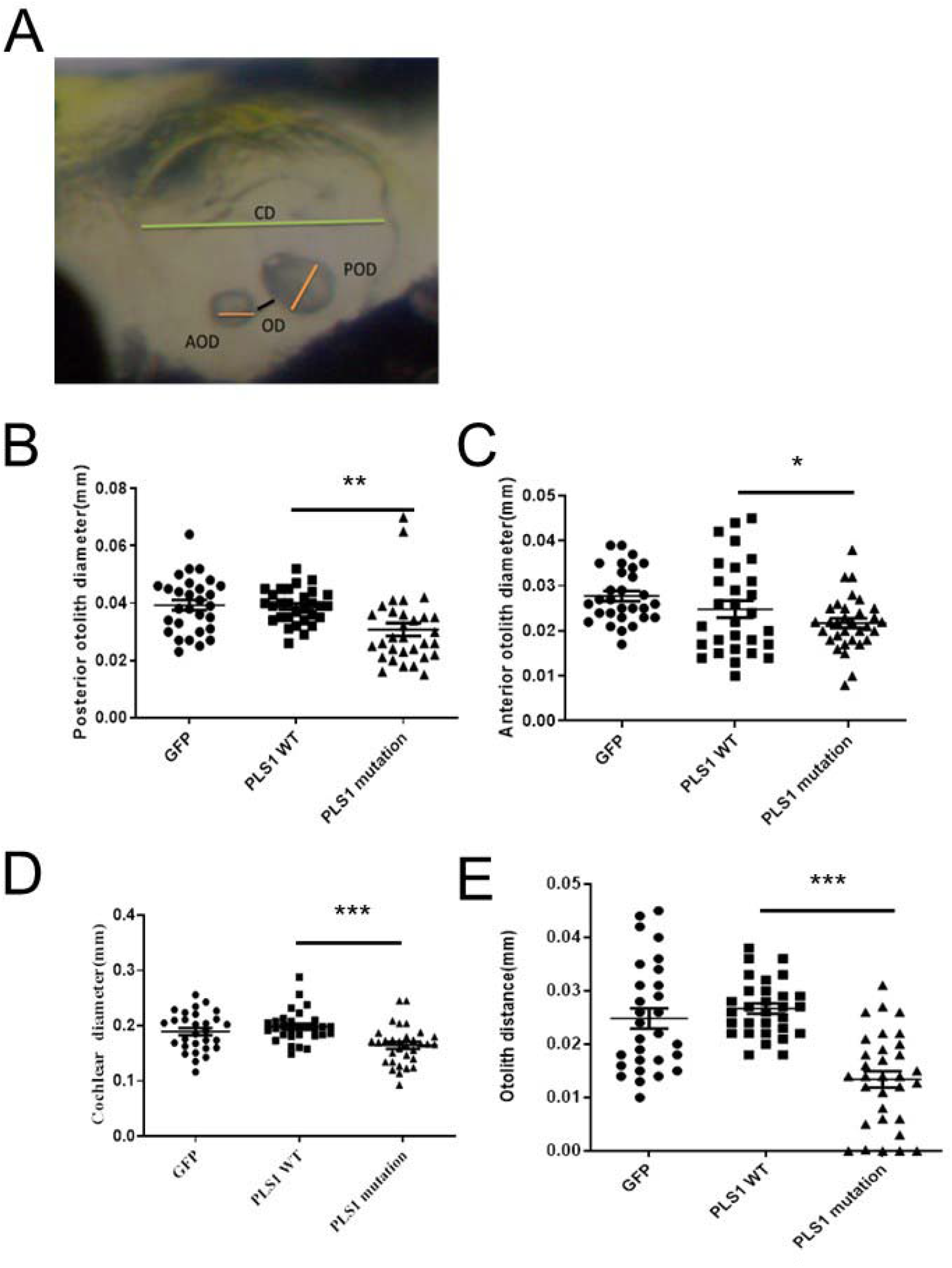
Comparison of otolith distance, anterior otolith diameter, posterior otolith diameter, and cochlear diameter in the 72 hpf zebrfish embryos injected with different mRNAs. (A) The diagram shows the measurement of zebrafish inner ear structures. OD, otolith distance; AOD, anterior otolith diameter; POD, posterior otolith diameter; AO, anterior otolith; CD, cochlear diameters. Comparison of (B) posterior otolith diameters, (C) anterior otolith diameters, (D) cochlear diameters, and (E) otolith spaces among the GFP (N=30), wildtype pls1 (N=30) and mutant pls1 zebrafish (N=30). WT: wildtype. *, **, *** indicate P < 0.05, P < 0.01, P < 0.001 respectively.

**Figure 4.**
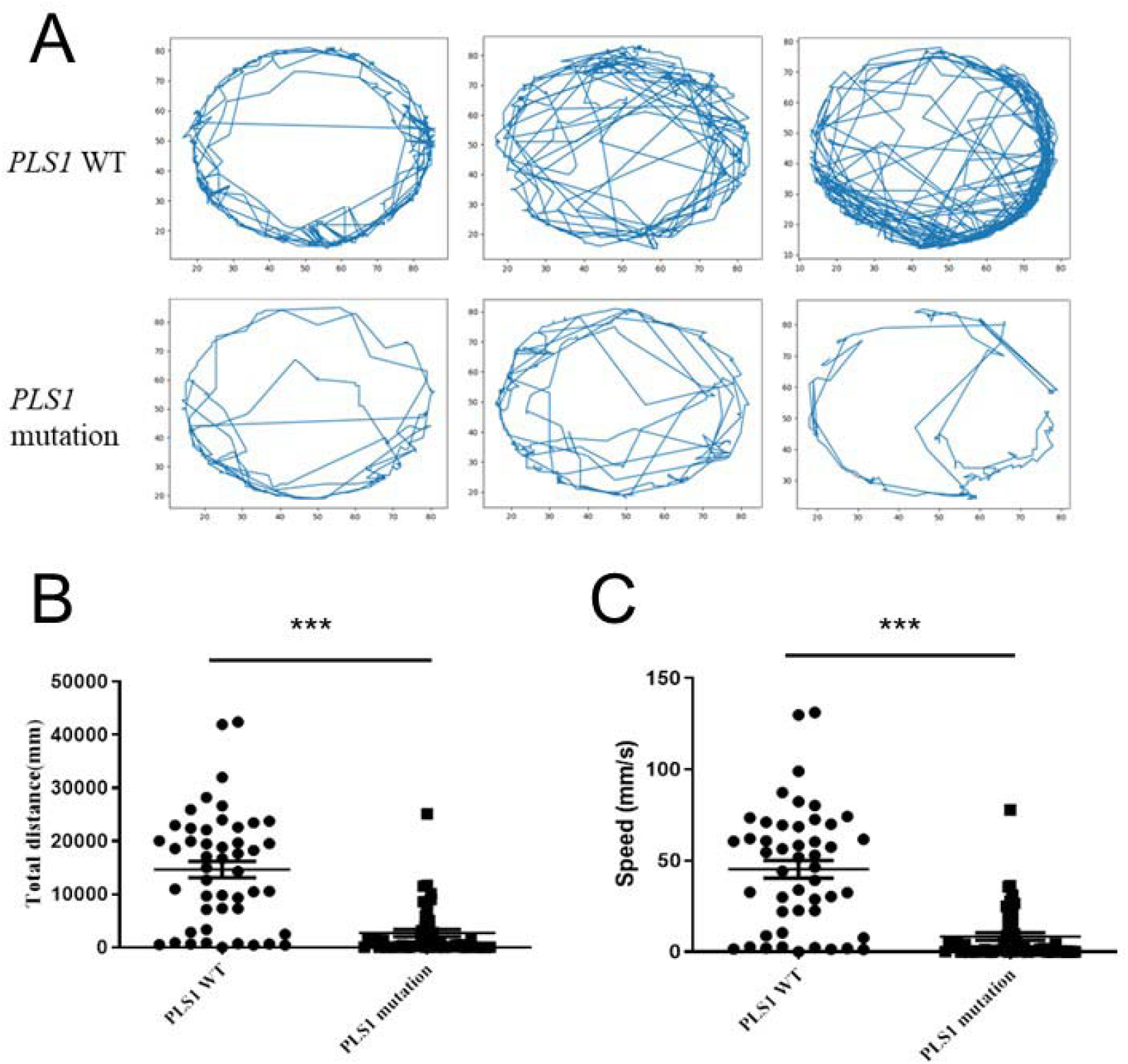
Results of total swimming distance and speed of wildtype pls1 and mutant pls1 zebrafish by zebrafish behavioral assay. (A) Swimming trajectory of the wildtype pls1 and mutant pls1 zebrafish. (B) Comparison of total swimming distance between the wildtype pls1 and mutant pls1 zebrafish. (C) Comparison of total swimming speed between the wildtype pls1 and mutant pls1 zebrafish. WT: wildtype, *** indicates P value <0.001.

### Pls-1 is involved in hearing loss via activating PI3K-Akt signaling pathway

To further investigate the mechanism by which Pls-1 is implicated in hearing loss, we transfected lentiviral shRNAs targeting Pls1 into HEI-OC1 cells and validated the knockdown efficiency using western blotting (Supplementary Figure6). Then, we performed transcriptional profiling on HEI-OC1 cells using RNA sequencing (RNA-seq) analysis. The results showed 478 genes were upregulated and 309 genes were downregulated in the shRNA treatment group (Figure 5A and Supplementary Figure7). Furthermore, the up-regulated genes were significantly enriched in signaling pathways, for example, focal adhesion, PI3K-Akt signaling pathway, ECM-receptor interaction (Figure 5B). Additional GO terms and Kyoto Encyclopedia of Genes and Genomes (KEGG) signaling pathways enrichment analysis of downregulated genes was shown in Supplemental Figure8. Furthermore, the up-regulated genes of the PI3K-Akt signaling pathway were validated by qPCR. The results confirmed significantly upregulated genes including Col6a3, Spp1, Itgb3 and Hgf in the shRNA treatment group (Figure 5C; P < 0.01 for all cases, Student t test). These results demonstrate that Pls-1 is implicated in hearing loss partially via regulating PI3K-Akt signaling pathway.

**Figure 5.**
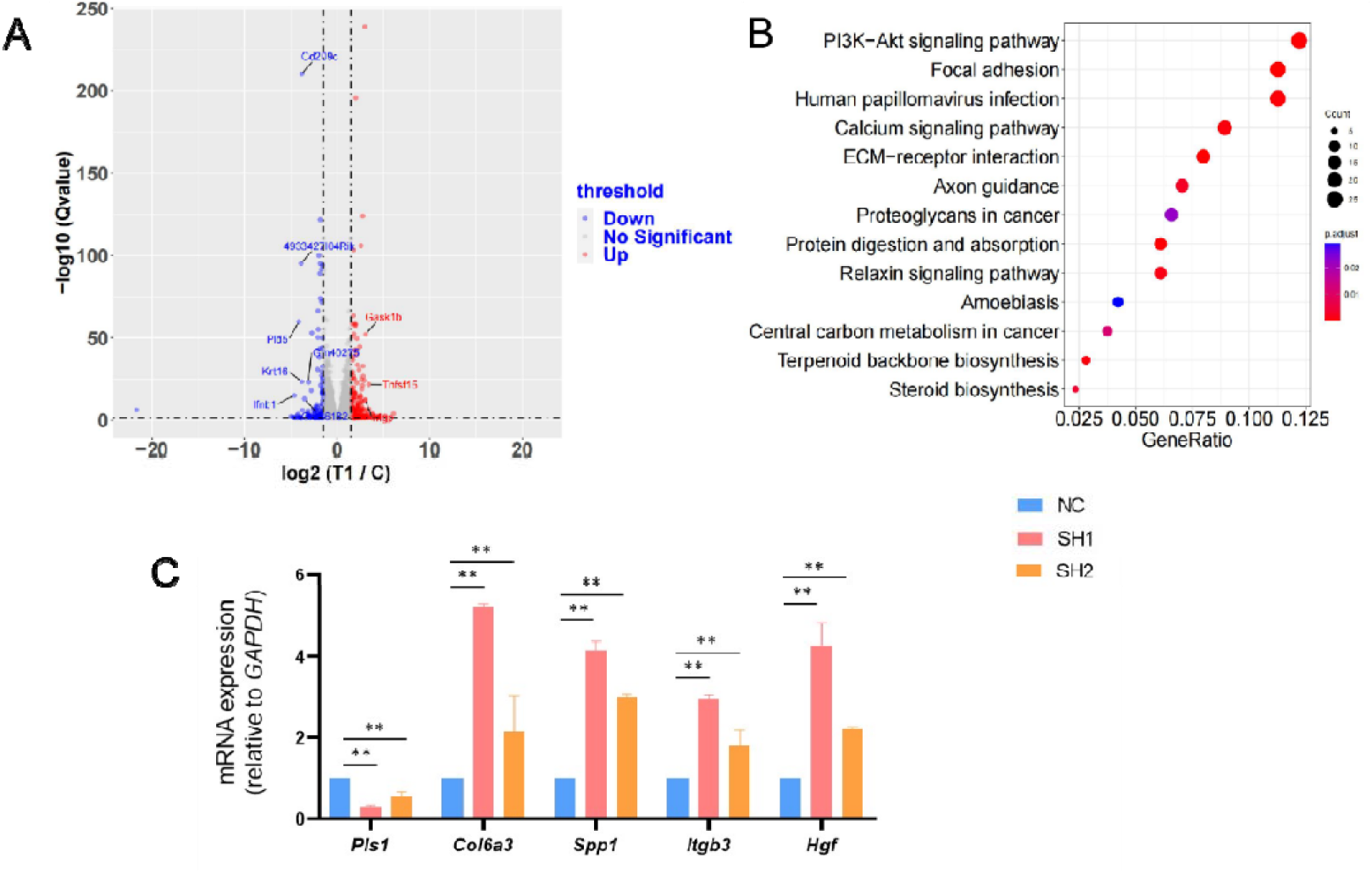
Characterization of dysregulated signaling pathways after silencing Pls-1 expression in HEI-OC1 cells. (A) Volcano plot showed differentially expressed genes in HEI-OC1 cells. Upregulated genes were shown in red, while downregulated genes in blue. **(B)** The KEGG signaling pathways significantly enriched for upregulated genes. **(C)** qPCR analysis confirmed knockdown of *PLS-1* led to the up-regulation of the PI3K-Akt signaling pathway genes in HEI-OC1 cells. NC: negative control, SH1: HEI-OC1 cells transfected with lentiviral shRNA 1 targeting Pls-1, SH2: HEI-OC1 cells transfected with lentiviral shRNA 2 targeting Pls-1. Data were shown as means ±standard deviation, *\**\* indicates *p* < *0*.*01*.

## Discussion

The PLS1 protein as one member of the human plastin family noncovalently crosslinks actin filaments into tight bundles^29^. Fimbrin plays an important role in the growth, integrity and function of stereocilia in mice^30^. Knockout of Pls1 in mice reduces the length and thickness of stereocilia, transforms actin filament packing from hexagonal to liquid, leading to a moderate and progressive hearing loss^12,13^. Taylor et al speculated that human PLS1 gene mutations may be related to mild and progressive forms of deafness. Several recent studies have reported the associations between mutations in the coding region of *PLS1* gene and the occurrence of deafness^14-16^. In this study, we identified a novel splicing variant in *PLS1* in one family affected by autosomal dominant NSHL. WES and sanger sequencing confirmed the variant cosegregated with hearing loss phenotype in the family, further minigene assay demonstrated the *PLS1* c.981+1G>A variant led to Exon8 skipping or deletion of Exon8 (47bp), suggesting that *PLS1* is required for normal hearing in human and mice.

In this study, we showed *PLS1* c.981+1G>A variant led to disrupted mRNA splicing and skipping of Exon 8 or deletion of Exon8 (47bp) [Amino acid: 297-327]. Furthermore, microinjection of mutant Pls1 mRNA into zebrafish embryos caused decreased mean otolith distance, anterior otolith diameter, posterior otolith diameter, and cochlear diameter, shorter swimming distance and lower swimming speed. The plastin protein family has a N-terminal regulatory region of 100 amino acids, including two EF-hand domains and two actin binding domains, which are composed of ABD1 (120-379) and ABD2 (394-623), respectively. Each ABD contains two calponin homologous (CH) domains^31^. ABD consists of a pair of calcium reducing protein homology (CH) domains with about 125 residues. Two ABD tandem repeat polypeptide chains of tow proteins and cross-linked actin filaments form higher-order assemblies through this pair of tandem ABDS^32^. The *PLS1* splicing variant identified is located in ABD.ABD1 binds one actin monomer in the filament and comprises two adjacent CH domains. Previous studies^33^ highlighted the two CH domains are essential for the actin-binding function. The *PLS1* c.981+1G>A variant led to disrupted mRNA splicing and skipping of Exon 8 or deletion of Exon8 (47bp) [amino acid: 297-327]. Therefore, we speculate that the disruption of the structure of ABD1 and its binding to actin may eventually lead to the failure of normal formation of stereocilia, resulting in hearing defects in all patients.

Though the pathogenic role of *PLS1* c.981+1G>A variant has been illustrated, the molecular mechanism by which PLS-1 is implicated in hearing loss remains unclear. Using RNA-seq analysis, we demonstrate that the up-regulated genes were significantly enriched in the PI3K-Akt signaling pathway and four up-regulated genes Col6a3, Spp1, Itgb3 and Hgf were further confirmed by qPCR. The PI3K/Akt signaling pathway play an important role in mediating survival of sensory hair cells and in the proliferation of auricular precursor cells^34,35^. Endogenous PI3K/Akt signaling has a protective role in the inner ear. Blockade of PI3K/Akt signaling pathways increases sensitivity to temporary-threshold-shift-inducing noise-induced hearing loss^36^. Of the up-regulated genes, hepatocyte growth factor (Hgf) is an activator of mitosis, stimulating epithelial cells to disperse in culture^37^ and plays an important role in normal hearing in human and mouse. A conditional Hgf deficiency in the ear may lead to thinning of the stria vascularis in the cochlea and viable deafness in mice^38^. Moreover, overexpression of HGF causes deafness in a *Hgf* transgenic mouse, suggesting that normal cochlear development is highly related to Hgf expression level^38,39^. Therefore, we believe the deafness caused by deficiency of Pls-1 may partially attributable to the up-regulation of PI3K/Akt signaling, particularly, Hgf gene.

## Fundings

This study was financially supported by Key Projects of Fujian Provincial Department of Science and Technology (Grant No. 2021J02054), Key Project on the Integration of Industry, Education and Research Collaborative Innovation of Fujian Province (Grant No. 2021YZ034011), Key Project on Science and Technology Program of Fujian Health Commission (Grant No. 2021ZD01002).

## Data available

The data reported in this article also available in the CNGB Nucleotide Sequence Archive(CNSA: https://db.cngb.org/cnsa/; accession number CNP0002720)

## Notes

### Competing Interest Statement

The authors have declared no competing interest.

