## supplemental materials for "A novel PLS1 c.981+1G>A variant causes autosomal-dominant hereditary hearing loss in a family via up-regulation of the PI3K-Akt signaling pathway"

**Supplemental Method 1. RNA-sequencing analysis**

**Library preparation**

Total RNA was isolated from the 9 samples, which three samples in each group, and RNA quality was assessed using an Agilent 2100 Bioanalyzer (Agilent Technologies, California, USA). The RNA was sheared and reverse transcribed using random primers to obtain cDNA which was used for library construction. The library was subsequently sequenced through BGISEQ-500 platform following a strategy of PE150 (MGI, Shenzhen, China) ([www.genomics.org.cn](http://www.genomics.org.cn/), BGI).

**Bioinformatics analysis**

Paired-end sequence files (fastq) were mapped to the reference genome (Mus_musculus, GCF_000001635.26_GRCm38.p6) using Hisat2 (Hierarchical Indexing for Spliced Alignment of Transcripts, version 2.0.4). The output SAM (sequencing alignment/map) files were converted to BAM (binary alignment/map) files and sorted using SAMtools (version 1.3.1). Gene abundance was expressed as fragments per kilobase of exon per million reads mapped (FPKM). Differential expression analysis for mRNA was performed using R package edgeR v3.18.1. Differentially expressed RNAs with |log2(FC)| value > 1 and adjusted P value < 0.05, considered as significantly modulated, were retained for further analysis. Potential gene function was analyzed based on enrichment of GO terms analysis for biological processes, cellular components and molecular function, and KEGG pathways analysis using the R package clusterProfiler v3.4.4.

Supplementary Table1. the primers for constructing the minigene vectors and for detecting alternative splice sites.

| **Primer** | **Sequences of primers** |
| --- | --- |
| pcMINI-N-PLS1-KpnI-F | GCTTGGTACCATGGACTCGAGAGCCTATTTTCA |
| pcMINI-N-PLS1-BamHI-R | TAGTGGATCCACAATTCTTCTTCTAATGTG |
| pcMINI-PLS1-KpnI-F | GGTAGGTACCTTCATCCTAGAGAATCCAGGGA |
| pcMINI-PLS1-BamHI-R | TAGTGGATCCAGCCTTGGGCATGGAAAAAAC |

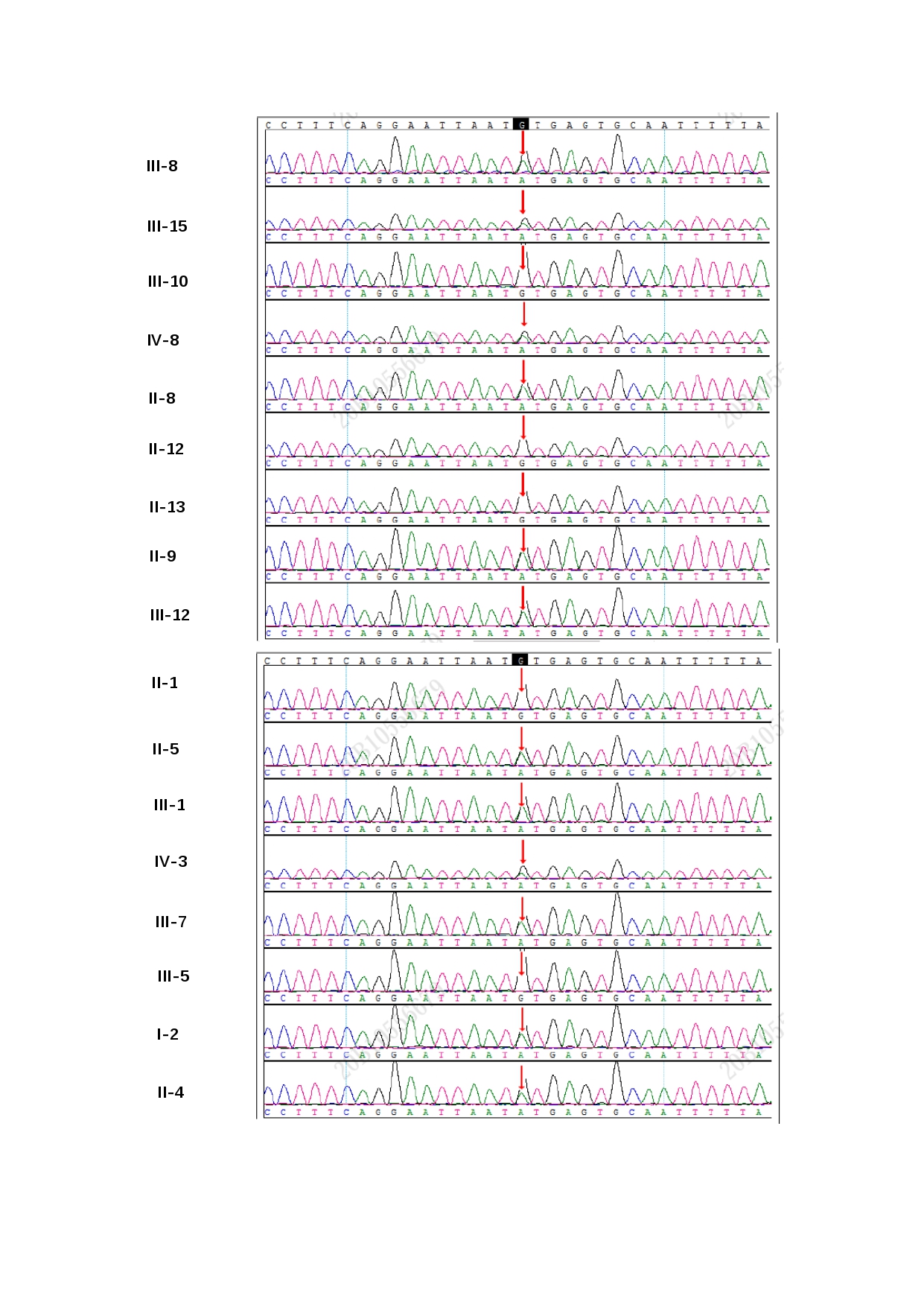

Supplementary Figure1. Sanger sequencing confirms the presence of heterogeneous *PLS1* c.981+1G>A in affected family members and absence of the variant in the healthy ones.

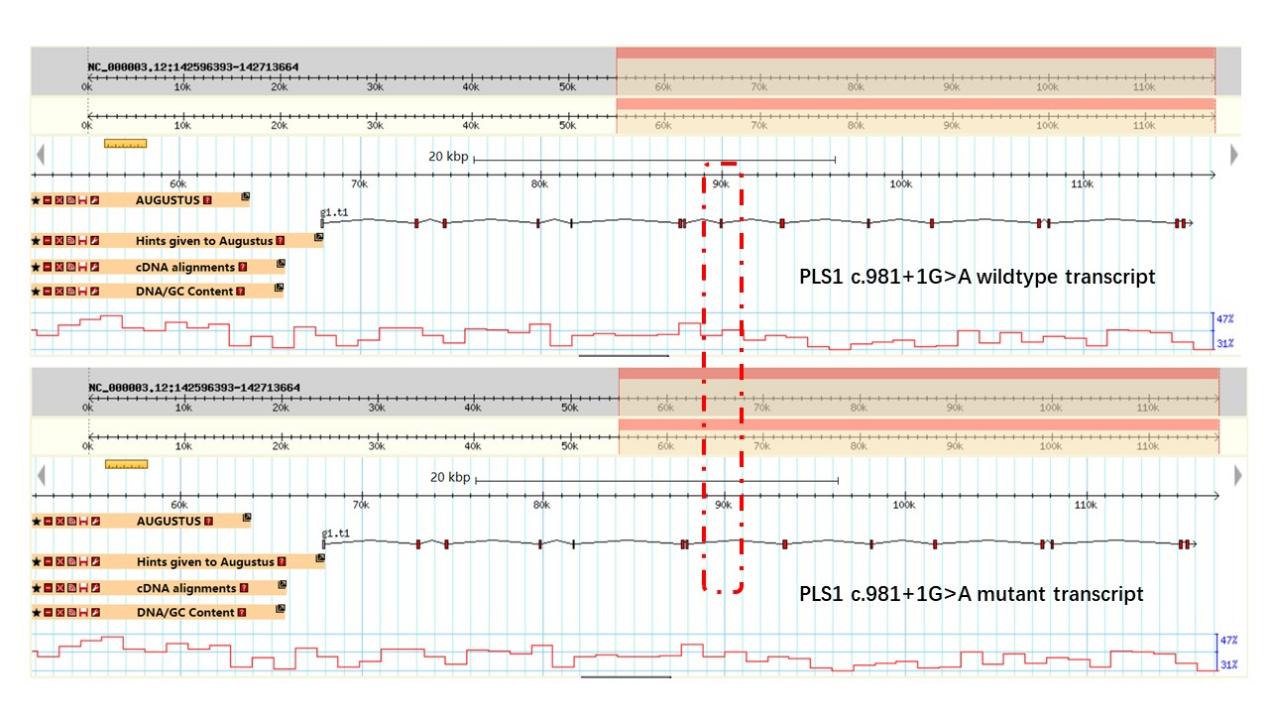

Supplementary Figure2. *PLS1* c.981+1G>A variant leads to Exon8 skipping predicted by the software augutus. The upper panel shows the *PLS1* c.981+1G>A wildtype transcript, while the lower panel shows the *PLS1* c.981+1G>A mutant transcript.

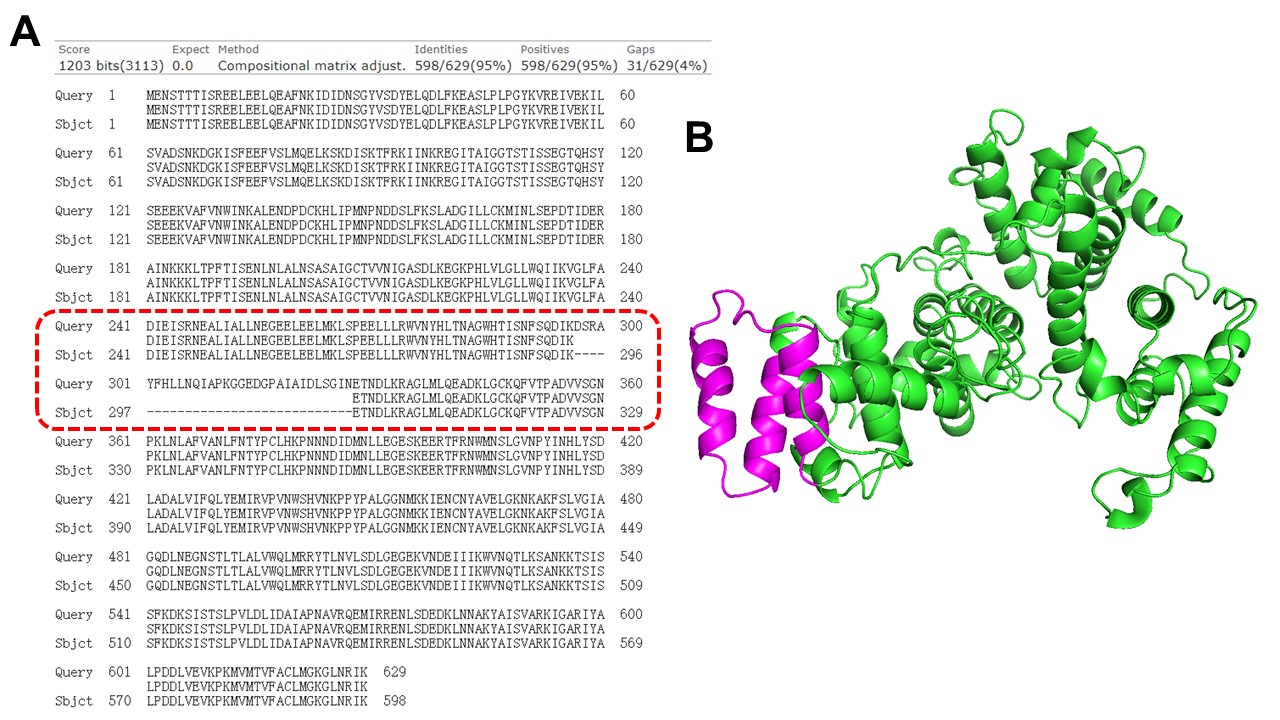

### Supplementary Figure3. Comparison of the 3D structure of mutant PLS1 and wild-type proteins. A. Comparison of amino acids between mutant PLS1 and wild-type proteins. Query: wildtype PLS1 amino acids, Sbjct: Mutant PLS1 amino acids. Deleted amino acids are shown in purple in the mutant PLS1 protein.

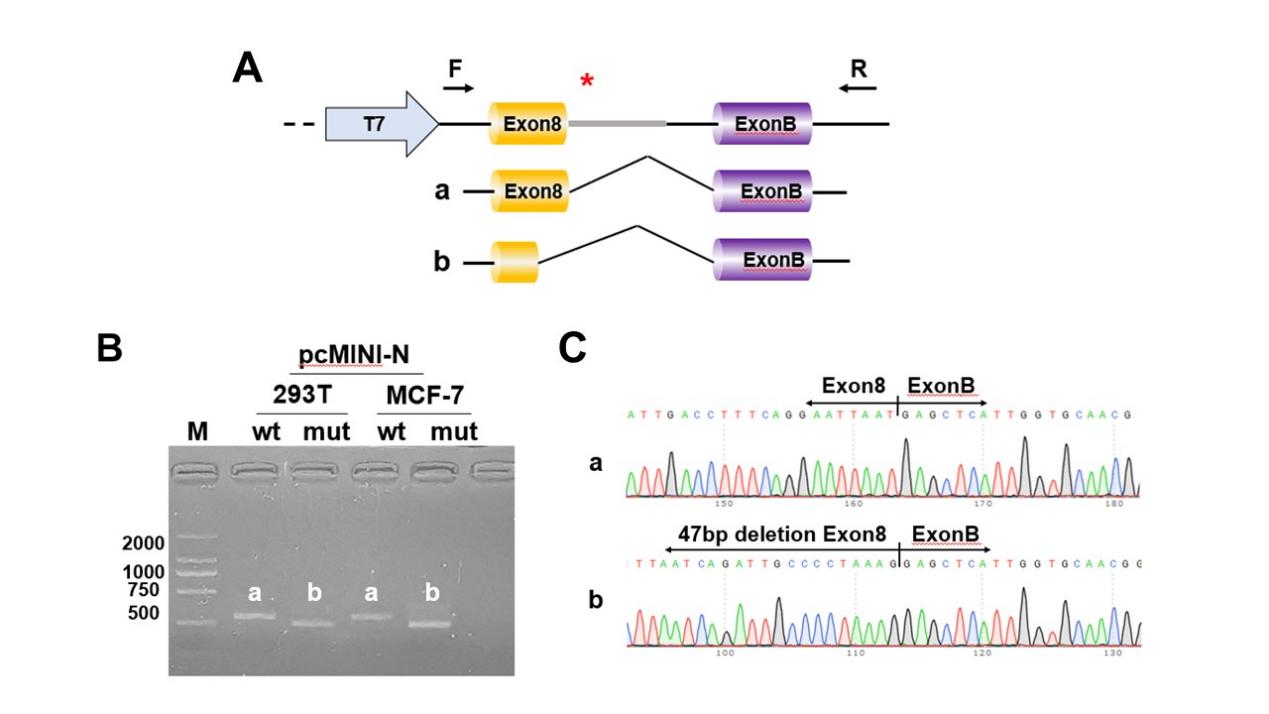

Supplementary Figure4. Results of minigene assay for *PLS1* c.981+1G>A in the HEK293T and MCF7 cell lines. A: The entire Exon8(93bp) and intron 8(798bp) of wild-type and mutant PLS1 were cloned into a report vector, pcMINI-N and transfected into HEK293T and MCF7 human cell lines. B. RT-PCR results of pcMINI-N- PLS1-mut and pcMINI-N- PLS1-wt constructs showed different sizes of bands in HEK293T and MCF-7 cells; C. Sanger sequencing demonstrated that band a presented a normal splicing pattern, while band b had 47 bp deletion of exon8.

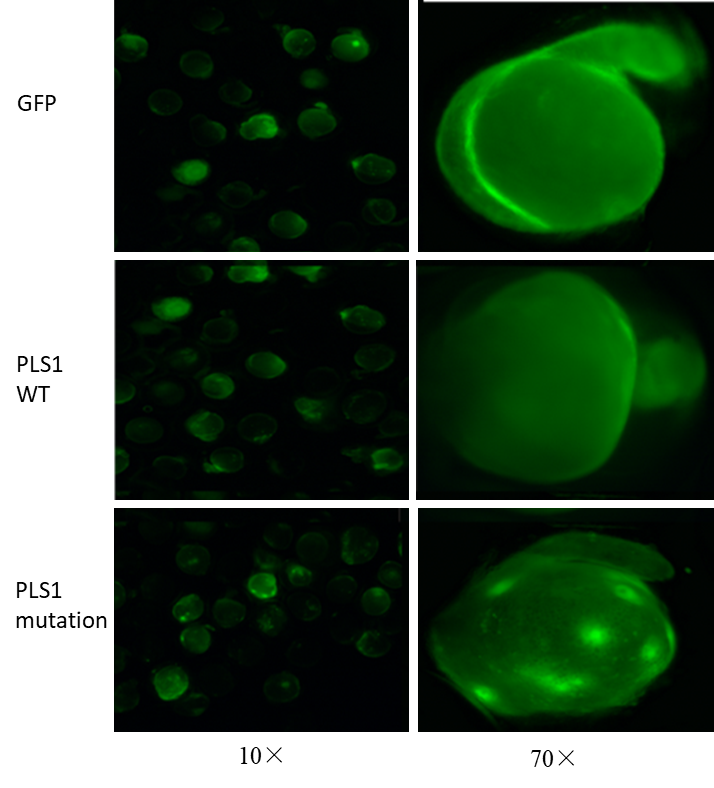

Supplementary Figure5. GFP was expressed in the 24 hpf zebrfish embryos injected with GFP mRNA, mutant PLS1 mRNA or wildtype PLS1 mRNA. WT: wildtype.

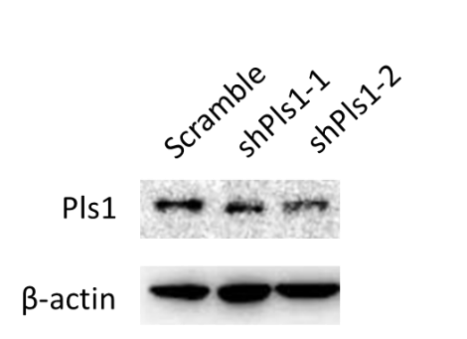

Supplementary Figure6. Comparison of PLS1 protein expression between scrambler (negative control), Pls1-knockdown group by lentiviral shRNA targeting Pls1.

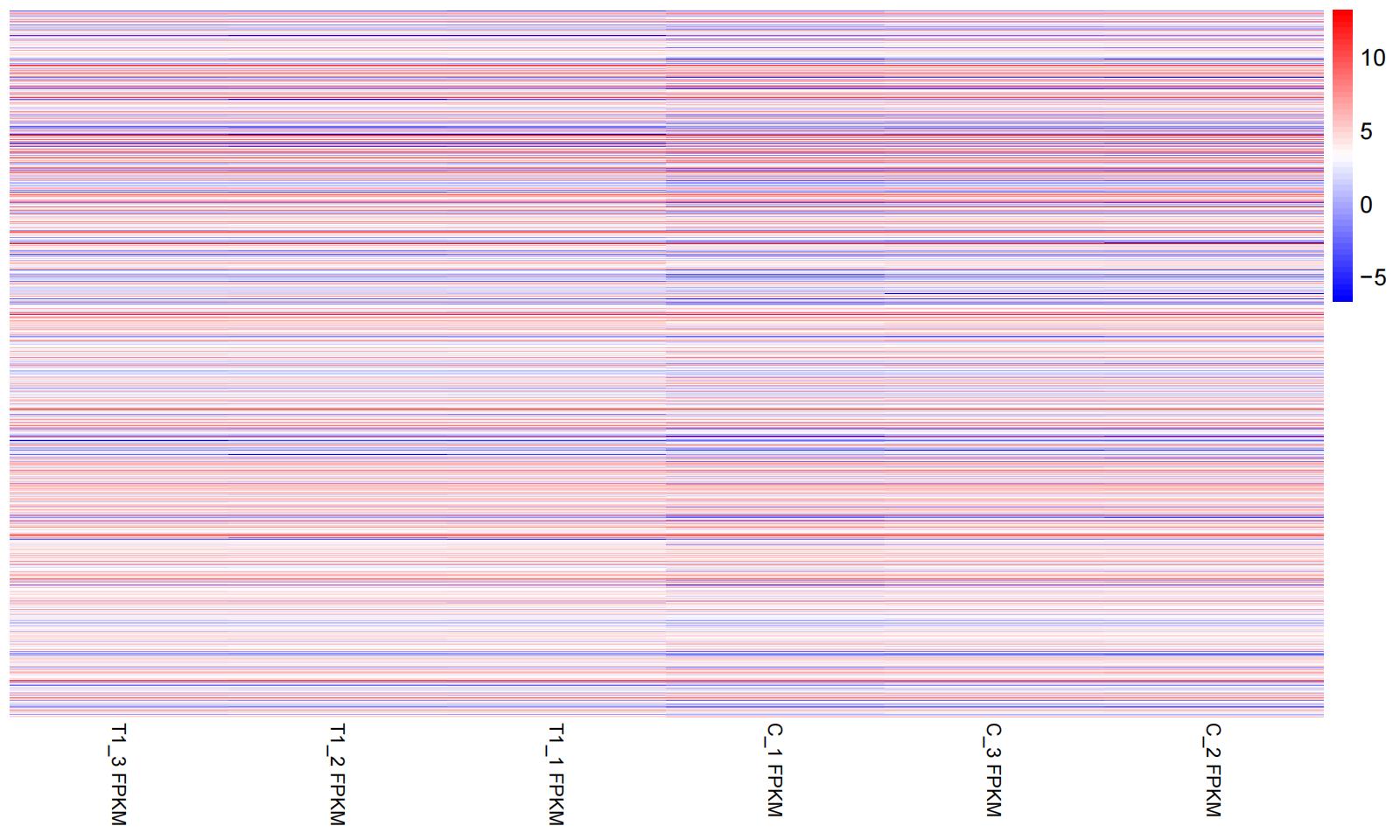

Supplementary Figure7.Heatmap of differentially expressed genes. T1_3, T1_2, T1_1 represent HEI-OC1 cells transfected with siRNA-1 targeting Pls1. C_1, C_2, C_3 denote negative controls.

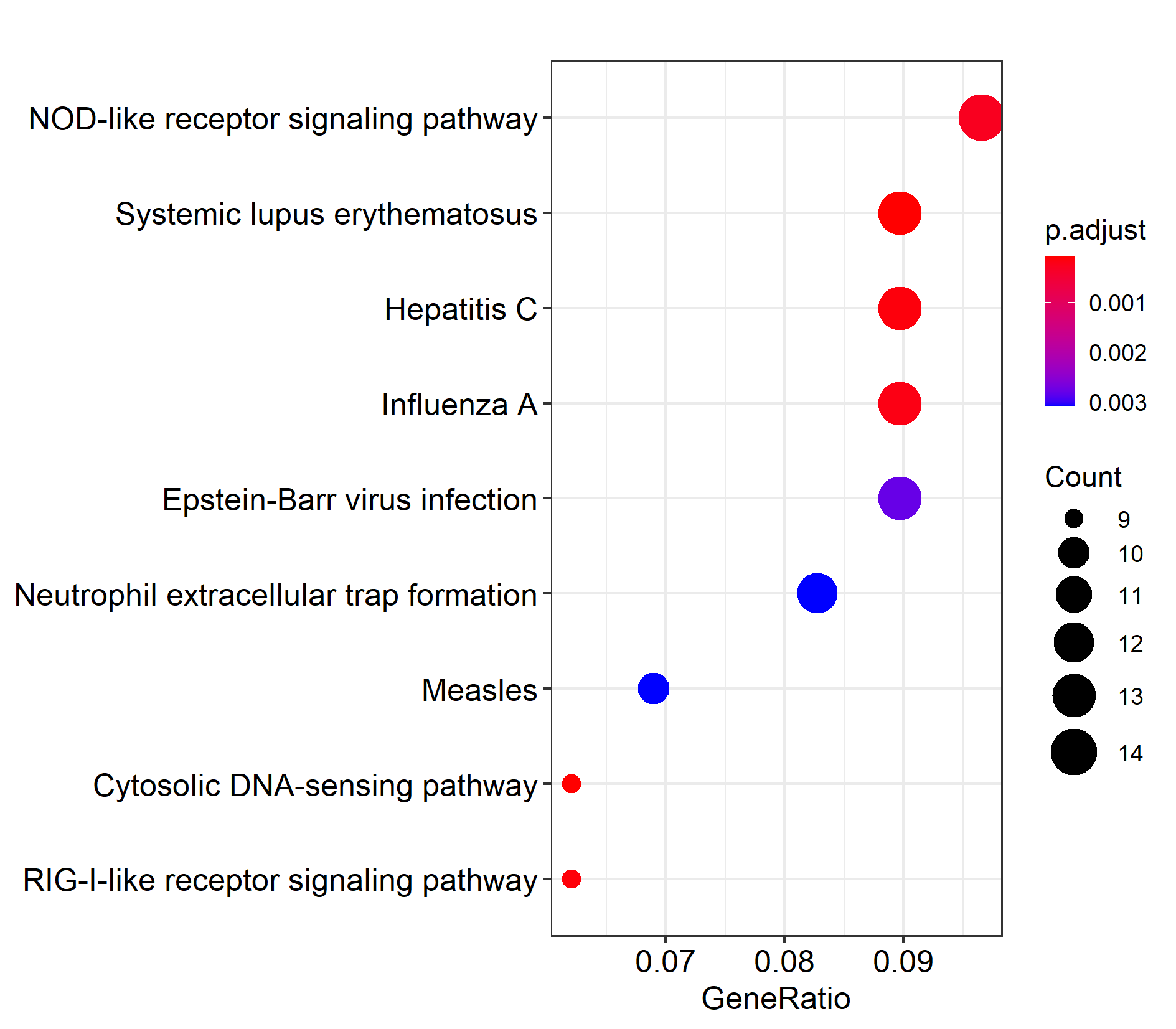

Supplementary Figure8. The significant KEGG signaling pathways enriched for down-regulated genes
